# Competition between protein-RNA clustering and phase separation drives re-entrant phase behavior of hnRNPA1

**DOI:** 10.1101/2025.07.25.666805

**Authors:** Katarzyna Makasewicz, Chiara Morelli, Lenka Faltova, Paolo Arosio

## Abstract

Phase separation of RNA-binding proteins plays crucial roles in the cell and is modulated by RNA, including promotion and suppression at low and high RNA concentrations, respectively. In complex coacervates, suppression of phase separation is rationalized by charge inversion when increasing the concentration of one component. Here, we show that suppression of biomolecular condensates of the RNA-binding protein hnRNPA1 at high RNA concentration is driven by a different mechanism, namely the competition with formation of nano-sized protein-RNA clusters in the dilute phase. We show that the competition is modulated not only by RNA concentration, but also by the type of RNA, with specific RNA being more effective in promoting cluster formation than unspecific RNA. We further show that protein-RNA clusters convert into amyloid fibrils over a longer time-scale compared to condensates, therefore providing higher kinetic stability. The competition between clustering and phase separation reported in this study provides a unifying framework to understand the distinct assemblies of hnRNPA1 in the nucleus and the cytoplasm, where the protein is exposed to different types and concentrations of RNA.

## Introduction

Phase separation of biomacromolecules in cells can lead to the formation of membraneless organelles, also known as biomolecular condensates.^1^ This process has been associated with a wide range of biological functions, including stress response, regulation of gene expression, and signaling.^2^ Cellular membraneless organelles are dynamic, and some form and dissolve in response to specific stimuli.^3^ Among other physico-chemical parameters, the formation of biomolecular condensates depends on the concentrations of their multiple components. Phase separation of RNA-binding proteins (RBPs) is modulated by RNA concentration.^4–7^ The effect of RNA was shown to be biphasic, with promotion and suppression of phase separation occurring at low and high RNA concentration, respectively, which has been referred to as re-entrant behavior. This phenomenon was proposed to have a regulatory role, wherein the cell can tune the formation and dissolution of liquid-like condensates through RNA levels.^4,6^

In systems of model peptides rich in RGG (arginine-glycine-glycine) motifs and oligonucleotides, reentrant behavior was shown to be driven by electrostatic interactions and modulated by short-range attractive interactions.^8,9^ Low concentrations of RNA drive peptide phase separation through charge neutralization induced by RNA binding, while the strength of short-range attractions between peptides and RNA governs the width of the two-phase regime.^9,10^ However, further increase in RNA concentration leads to suppression of phase separation due to overscreening of the peptide macromolecular charges by RNA and long-range electrostatic repulsion between the overcharged peptides.

However, biological systems displaying re-entrant phase behavior feature proteins with complex architectures including both disordered and folded domains with different propensities for self- and co-assembly, which modulate the solubility of the protein^11^ and engage in both sequence-specific and unspecific interactions with RNA.^12,13^ Molecular mechanisms of re-entrant phase separation of full length RNA-binding proteins have not been studied before.

In this work, we studied the molecular mechanism of suppression of hnRNPA1 phase separation at high RNA concentration. hnRNPA1 is a member of the family of heterogeneous nuclear ribonuclear proteins (hnRNPs). Its domain architecture, characteristic of many RNA-binding proteins, consists of a folded domain and a disordered low complexity domain (LCD) containing RGG repeats and a prion-like domain.^14^ The folded domain of hnRNPA1, known as unwinding protein 1 (UP1), features two RNA Recognition Motifs (RRMs), which engage in sequence-specific RNA binding,^15^ and modulate the solubility of the protein.^11^ UP1 does not undergo phase separation on its own, while the low complexity domain is necessary and sufficient for phase separation.^11,16,17^

*In vivo*, hnRNPA1 is present both in the nucleus and in the cytoplasm, with its concentration in the nucleus being approx. 3 times higher than in the cytoplasm.^4,18,19^ In the nucleus, the protein plays a role in transcriptional regulation, associates with pre-mRNA, acts as a splicing factor and takes part in 3’ end processing.^20,21^ hnRNPA1 shuttles at a high rate between the nucleus and the cytoplasm and plays a role in nucleocytoplasmic transport of mRNA.^18,19^ hnRNPA1 does not undergo phase separation in the nucleus, which was hypothesized to be due to high RNA concentration,^4^ consistent with its phase separation being enhanced at low RNA concentration and suppressed at high RNA concentration.^4,5^ Under stress conditions, hnRNPA1 is transported to the cytoplasm where it is sequestered in stress granules.^22^

Here, we show that suppression of hnRNPA1 phase separation at high RNA concentration is driven by the competition with the formation of nano-sized protein-RNA clusters in the dilute phase. We provide evidence that this competition is mediated by the multi-domain structure of hnRNPA1, highlighting the interplay of folded and unstructured domains in modulating protein phase behavior. We show that the competition is modulated not only by RNA concentration, but also by the strength of protein-RNA interactions, with specific RNA being more effective in promoting cluster formation than unspecific RNA, although their effect in inducing condensates at low concentrations is similar. Our results demonstrate that dissolution of RBP condensates at high RNA concentration is driven by a fundamentally different mechanism compared to systems of oppositely charged polymers, although the macroscopic behavior is similar, and show how cells can exploit competition between co-assembly and phase separation to dynamically modulate protein phase behavior.

## Results

### Biphasic effect of RNA on phase separation of hnRNPA1

First, we investigated the phase separation of hnRNPA1 in the presence of increasing concentrations of RNA. The RNA used in this study is a single-stranded 18 nt long sequence (5*^′^*-CCA GCA UUA UGA AAG UGA-3*^′^*) derived from human intronic splicing silencer N1 from the SMN2 gene transcript (SMN2-ISS-N1), which was previously shown to bind with high affinity to RRMs of hnRNPA1.^15,23^ We acquired confocal fluorescence images of samples containing 10 *µ*M hnRNPA1 (incl. 500 nM hnRNPA1 labelled with Atto 647) and increasing concentrations of RNA (up to 10 *µ*M RNA incl. 0.5 *µ*M RNA labelled with fluorescein (FAM)) (Figure 1A). Low concentrations of RNA (4 *µ*M and below) promote phase separation, as indicated by increased number and size of condensates compared to conditions without RNA. However, at 5 *µ*M RNA, the observed condensates are fewer and smaller, indicating that the driving force for phase separation decreases. At concentrations equal or larger than 10 *µ*M RNA, hnRNPA1 phase separation is suppressed. The biphasic effect of RNA on hnRNPA1 phase separation was further confirmed by the measured fluorescence intensity in the dilute phase, which reports on the concentrations of hnRNPA1-647 and RNA-FAM (Figure 1B), as well as by turbidity (right angle scattering) measurements, reporting on the amount of the dense phase (Figure 1C).

**Figure 1:**
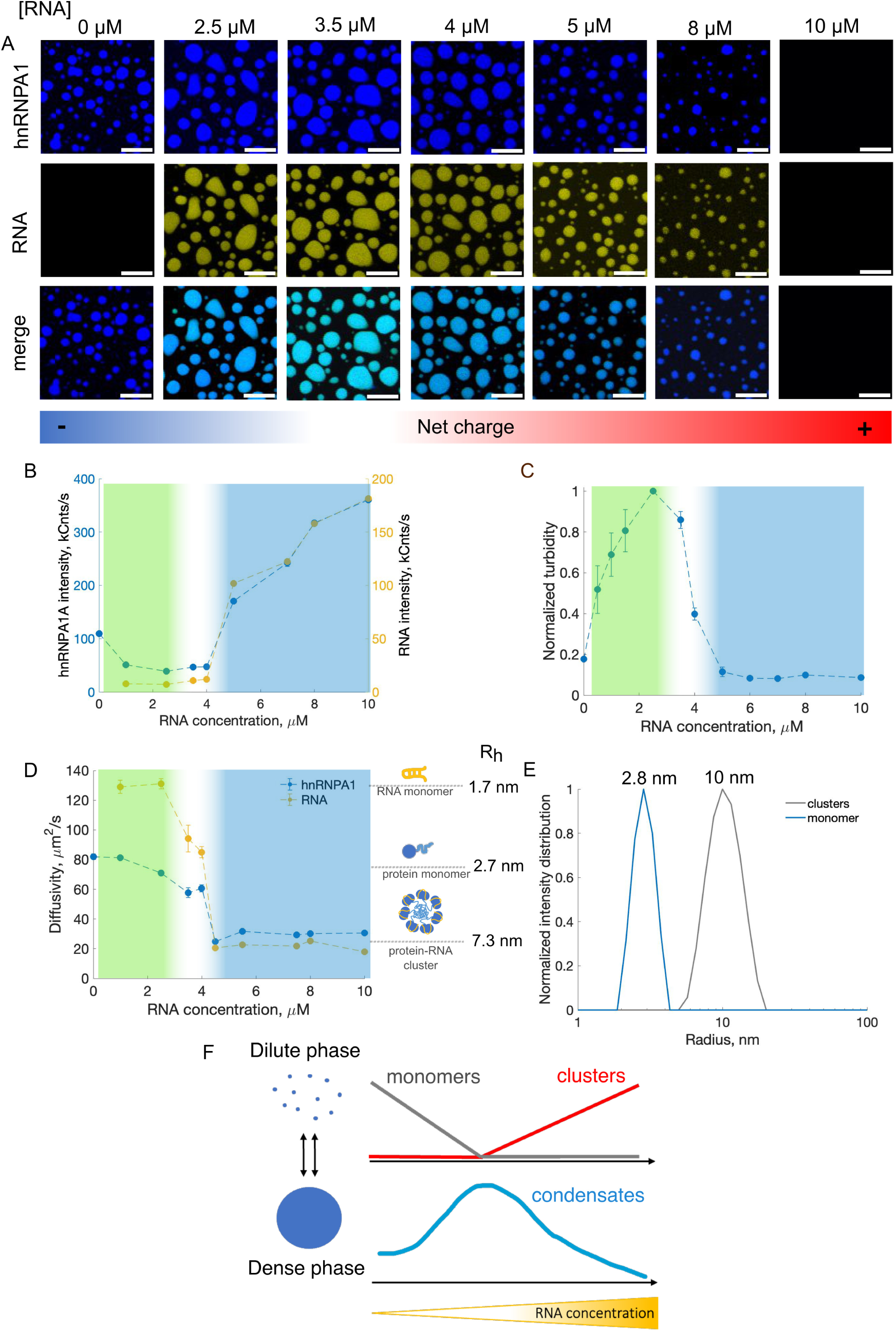
Suppression of hnRNPA1 phase separation at high RNA concentrations is accompanied by protein-RNA co-assembly in the dilute phase. A) Confocal fluorescence images of samples containing 10 *µ*M hnRNPA1 (incl. 0.5 *µ*M hnRNPA1A-647) and 0-10 *µ*M RNA (incl. 0.5 *µ*M RNA-FAM). The scale bar in all images is 10 *µ*M. B) Fluorescence intensity of hnRNPA1-647 (blue) and RNA-FAM (yellow) measured using Fluorescence Correlation Spectroscopy (FCS) in the dilute phase of the samples shown in A. The error bars are smaller than data points. C) Normalized turbidity (right angle scattering) measured for samples containing 10 *µ*M hnRNPA1 and increasing concentrations of RNA. D) Diffusivity of hnRNPA1-647 and RNA-FAM measured in the dilute phase of samples shown in A using FCS. FCS autocorrelation curves are shown in Figure S1. Colored regions in panels B-D correspond to regimes where RNA promotes hnRNPA1 phase separation and increase in RNA concentration leads to an increase in the volume of the dense phase (green), where RNA suppresses hnRNPA1 phase separation (blue) and a transition regime where hnRNPA1 phase separation is promoted with respect to protein-only sample but increasing RNA concentration leads to a decrease in the volume of the dense phase (white). E) Dynamic light scattering of monomeric hnRNPA1 (10 *µ*M hnRNPA1 in 20 mM TRIS buffer with 500 mM NaCl) and of protein-RNA clusters formed in a sample containing 10 *µ*M hnRNPA1 and 10 *µ*M RNA. F) Characterization of the dilute phase coexisting with the condensates as a function of RNA concentration reveals a change in the identity of the dilute phase, from one containing protein and RNA monomers to one containing protein-RNA clusters, which occurs at the point of maximum propensity for phase separation.

### Competition between phase separation and protein-RNA clustering in the dilute phase

In order to gain insights into the molecular mechanism underlying suppression of hnRNPA1 phase separation at high RNA concentration, we analyzed the behavior of protein and RNA in the dilute phase that coexists with the condensates (Figure 1F). To this end, we employed Fluorescence Correlation Spectroscopy (FCS), which allows us to measure the diffusivity and concentration of fluorescently-labeled species (Figure 1B, D). We analyzed the same samples previously investigated by microscopy (Figure 1A).

We started by analyzing the diffusivity of hnRNPA1 (Figure 1D and Figure S1). At low RNA concentration (2.5 *µ*M and below), the diffusion coefficient extracted from the fitting of the autocorrelation curves is 80 ± 8 *µ*m^2^/s and corresponds to a hydrodynamic radius of 2.7 nm, indicating that hnRNPA1 in the dilute phase is monomeric (Figure S2). In the intermediate RNA concentration regime (3.5 and 4 *µ*M), hnRNPA1 diffusivity decreases to approx. 60 *µ*m^2^/s, which indicates that small protein assemblies emerge in the dilute phase. At RNA concentration of 5 *µ*M and above, the dilute phase consists of larger protein clusters with diffusion coefficient of 30 ± 3.7 *µ*m^2^/s, corresponding to a hydrodynamic radius of 7.3 nm. The macroscopically homogeneous phase present at high RNA concentration ([*RNA*] = 10 *µ*M) consists entirely of such clusters (see additional analysis shown in Figure S2). In addition to the diffusivity, we analyzed the fluorescence intensity of hnRNPA1-647 in the dilute phase, which reports on the protein concentration (Figure 1B). The initial increase of RNA concentration results in a decrease in protein concentration, consistent with promotion of phase separation, while a further increase of RNA concentration leads to an increase of protein concentration in the dilute phase, consistent with phase separation being suppressed. We note that at low RNA concentration, fluorescence intensity reports on monomeric hnRNPA1 concentration, while at high RNA concentration, it reports on the concentration of hnRNPA1 clusters.

We next employed FCS to analyze the diffusivity of RNA in the dilute phase. Changes in RNA diffusivity with increasing RNA concentration followed the same trend as observed with hnRNPA1 (Figure 1D), suggesting that the protein clusters contain also RNA. We confirmed this result using fluorescence cross-correlation spectroscopy (FCCS), which probes the dynamic co-localization of fluorescently-labeled molecules (Figure S3). FCCS data revealed the absence and presence of protein-RNA species in the dilute phase at low and high RNA concentration, respectively. Thus, the soluble clusters observed at high RNA concentration contain both hnRNPA1 and RNA.

Taken together, the diffusivity and turbidity data allow us to divide the studied RNA concentration range into three regimes: in the first regime, increasing RNA concentration promotes phase separation, and hnRNPA1 in the dilute phase is monomeric (green-shaded region in Figures 1B-D). The second regime starts at the point of maximum phase separation (2.5 *µ*M RNA concentration), where condensates are still present, but a further increase of RNA concentration reduces the volume of the dense phase. In this regime (white region in Figures 1B-D), protein-RNA clusters of intermediate size appear in the dilute phase. In the third regime (blue-shaded region in Figures 1B-D), the high RNA concentration suppresses hnRNPA1 phase separation, and the dilute phase consists of protein-RNA clusters. In this regime, the size of the clusters remains constant, but their amount increases as the volume of the dense phase decreases (Figure 1B and D). These results suggest that hnRNPA1 phase separation and soluble cluster formation are in competition, which is modulated by RNA concentration (Figure 1F and Figure 5A).

From the comparison of the diffusivity of the hnRNPA1 monomer and clusters, we estimated that the clusters contain on average 20 protein monomers (details in Supplementary Information). Moreover, based on FCS data we estimated that RNA:protein stoichiometry in the clusters formed at 10 *µ*M hnRNPA1 and 10 *µ*M RNA is approximately 1:1 (details in Supplementary Information). This is consistent with the absence of detected monomeric protein under these conditions (Figure S2) and implies that all protein and RNA are incorporated into the clusters.

We confirmed the FCS analysis with dynamic light scattering measurements on unlabelled protein and RNA molecules (Figure 1E). The results confirmed the presence of clusters with the size distribution centered at 10 nm in radius and no detectable monomeric protein in the one-phase sample at high RNA concentration.

### Effect of RNA chemistry

Next, we analyzed the effect of RNA chemistry on the hnRNPA1-RNA co-assembly by comparing the effects of the 18nt SMN2-ISS-N1 RNA and a model RNA oligomer U-20, which is not expected to have any specific interactions with hnRNPA1. As shown by confocal microscopy (Figure 2A,B) and turbidity measurements (Figure 2C), U-20 exerts a biphasic effect on hnRNPA1 phase separation similar to the behavior observed with RNA, but a higher concentration is needed to suppress phase separation.

**Figure 2:**
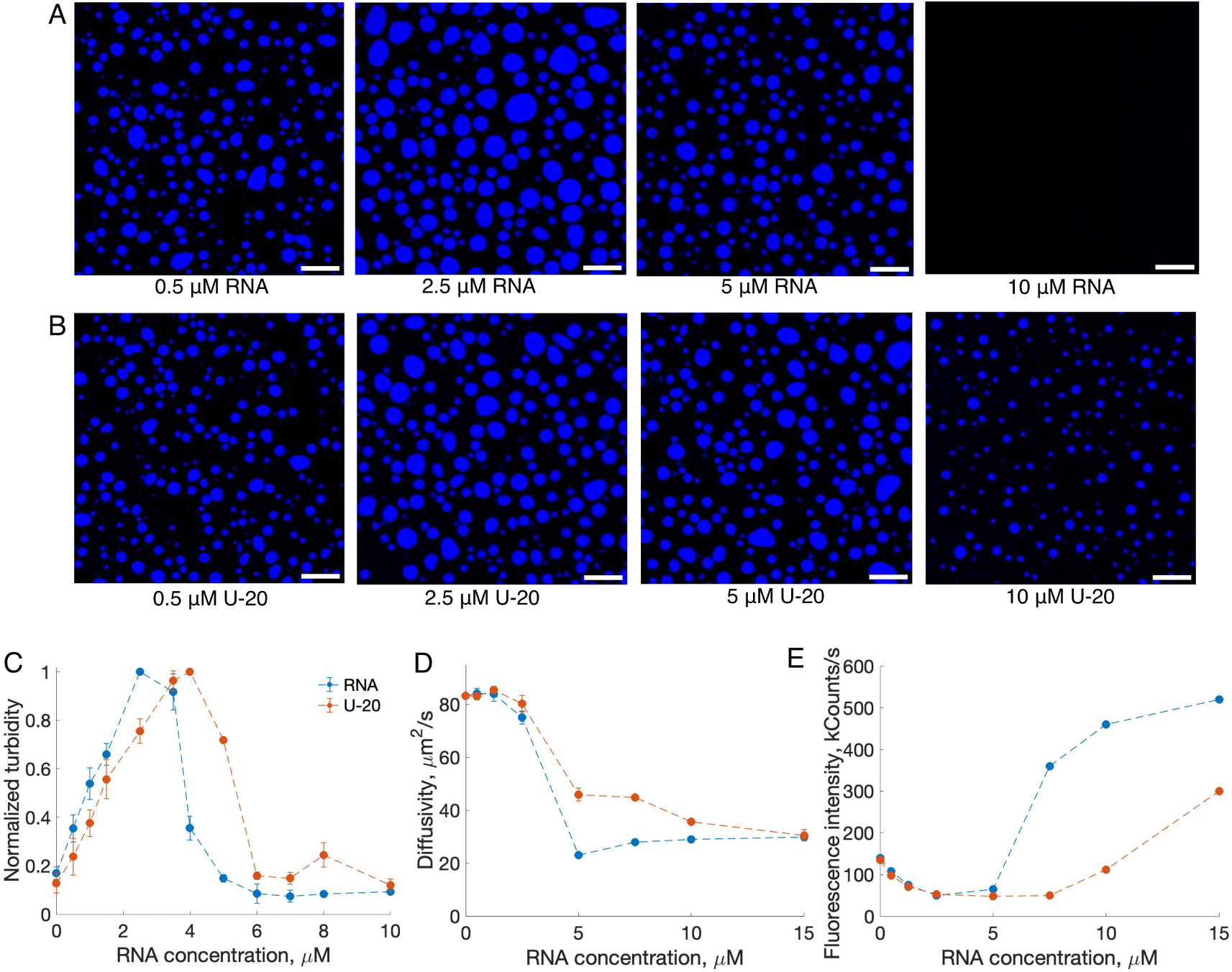
Effect of RNA chemistry on hnRNPA1 phase separation and protein-RNA clustering in the dilute phase. A) Confocal images of 10 *µ*M hnRNPA1 (incl. 0.5 *µ*M hnRNPA1A-647) with increasing concentration of RNA. B) Confocal images of 10 *µ*M hnRNPA1 (incl. 0.5 *µ*M hnRNPA1A-647) with increasing concentration of U-20. The scale bar is 10 *µ*m. C) Turbidity measured for samples containing 10 *µ*M hnRNPA1 and increasing concentrations of RNA/U-20. D) Diffusivity of hnRNPA1 in the dilute phase of samples containing 10 *µ*M hnRNPA1 and increasing concentrations of RNA/U-20. E) Fluorescence intensity of hnRNPA1-647 in the dilute phase measured with FCS for the same samples as analyzed in D. Error bars are smaller than data points.

We again employed FCS to analyze the diffusivity of the protein in the dilute phase of samples consisting of 10 *µ*M hnRNPA1 and increasing concentrations of U-20 and compared it with data acquired at the same concentrations of RNA (Figure 2D). To avoid any effects of the presence of fluorescent labels on RNA, we used 100% unlabelled RNA and U-20. In the case of RNA, we again observed a relatively sharp transition from monomeric protein to clusters with diffusivity of 30 *µ*m^2^/s at 5 *µ*M RNA and no further increase in cluster size with increasing RNA concentration. In the case of U-20, however, clusters of intermediate size with diffusivity of 45 ± 5 *µ*m^2^/s were present at 5 and 7.5 *µ*M U-20, while clusters with mean diffusivity of 30 *µ*m^2^/s emerged only at 15 *µ*M U-20. Comparison of the fluorescence intensity of hnRNPA1-647 in the dilute phase revealed no difference in the effect of RNA and U-20 at low concentrations, i.e. in the phase separation-promoting regime (Figure 2E). However, at higher RNA concentrations, i.e. in the phase separation-suppressing regime, lower intensity of hnRNPA1-647 was observed in presence of U-20 compared to RNA, indicating that less protein is incorporated in the clusters in the dilute phase at a given RNA/U-20 concentration.

These results indicate that U-20 is less effective in driving cluster formation and requires higher concentrations to suppress phase separation compared with RNA. Therefore, the efficiency of RNA in driving hnRNPA1 cluster formation correlates with the efficiency of suppression of phase separation, which supports the conclusion that clustering and phase separation are in competition (Figure 5A). Furthermore, these findings indicate that sequence-specific, high-affinity protein-RNA interactions play a more important role in suppressing than promoting hnRNPA1 phase separation.

### The role of hnRNPA1 multi-domain architecture in the formation of protein-RNA clusters

Next, we analyzed the molecular determinants underlying the observed competition between phase separation and cluster formation. We first compared the behavior of the full-length protein hnRNPA1 with the low-complexity domain alone (indicated in the following as A1-LCD). We analyzed samples containing 10 *µ*M A1-LCD and increasing concentrations of RNA. RNA promoted phase separation of A1-LCD, as shown by microscopy confocal images (Figure 3A) and by the decrease of fluorescence intensity of A1-LCD in the dilute phase, which is proportional to the protein concentration (Figure S4). However, in contrast to full length hnRNPA1, no suppression of phase separation was observed even at the highest RNA concentration tested (40 *µ*M). Furthermore, FCS diffusivity measurements in the dilute phase coexisting with condensates revealed that A1-LCD is monomeric under all tested conditions (Figure 3B, Figure S5A). Thus, the presence of the folded domain is required for the formation of hnRNPA1-RNA clusters and the suppression of phase separation at high RNA concentration.

**Figure 3:**
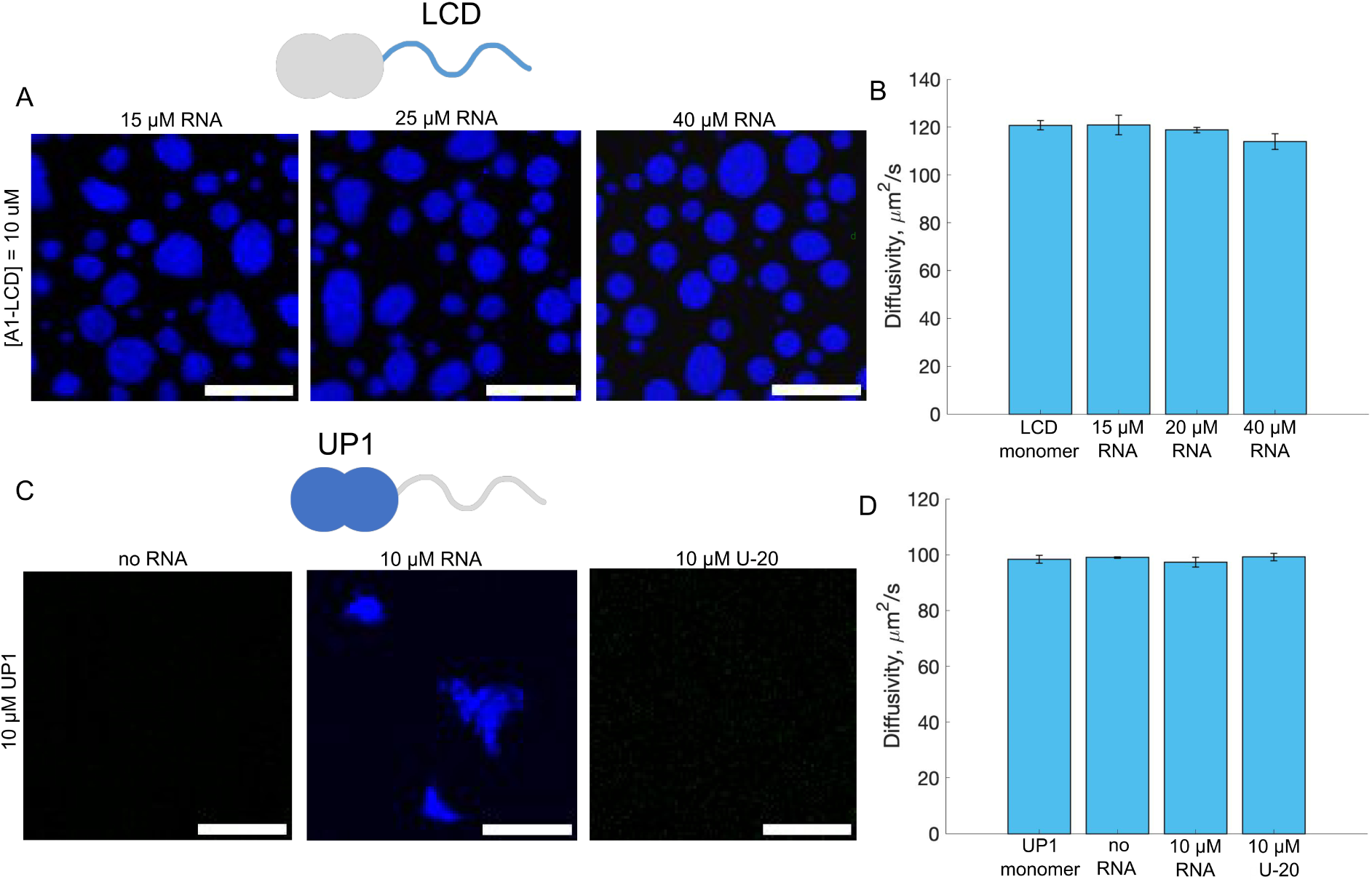
The role of multi-domain architecture of hnRNPA1 in protein-RNA clustering. A) Phase separation of A1-LCD is not suppressed at high RNA concentrations. Confocal images of 10 *µ*M A1-LCD (incl. 500 nM A1-LCD-647) with 15, 25 and 40 *µ*M RNA. Scale bar is 10 *µ*m. B) Analysis of the dilute phase coexisting with A1-LCD-RNA condensates. FCS diffusivity measurements show that A1-LCD is monomeric under all conditions tested. FCS autocorrelation curves are shown in Figure S5A. C) Folded domain of hnRNPA1 (UP1) does not undergo phase separation in absence (left) and in presence of U-20, while amorphous precipitates form in presence of specific RNA (middle). Confocal images of 10 *µ*M UP1 (incl. 500 nM UP1-565) in absence or presence of 10 *µ*M specific RNA and U-20. Scale bar is 10 *µ*m. D) Diffusivity measured in the dilute phase of samples shown in C shows that UP1 is monomeric under all conditions tested. FCS autocorrelation curves are shown in Figure S5B.

The folded domain of hnRNPA1 contains two RNA recognition motifs, which engage in sequence-specific interactions with RNA.^15,23^ The fact that RNA is more effective than U-20 in driving hnRNPA1 cluster formation suggests that the high-affinity sequence-specific interaction between the RRMs and RNA plays an important role. To investigate this further, we analyzed samples containing 10 *µ*M folded domain of hnRNPA1 (indicated as UP1 in the following) in the absence and presence of 10 *µ*M RNA or U-20. Consistent with the low-complexity domain being necessary for phase separation,^16^ confocal image of 10 *µ*M UP1 shows homogeneous fluorescence and an absence of condensates (Figure 3C). The diffusivity of UP1 measured in this sample equals 98.4 ± 2.7 *µ*m^2^/s and corresponds to diffusivity of monomeric UP1 measured in high salt buffer (Figure 3D, Figure S5B). Protein diffusivity in samples containing 10 *µ*M RNA or U-20 also corresponded to monomeric UP1, indicating no cluster formation in solution. The sample with U-20 was macroscopically homogeneous, while in the sample with RNA, amorphous precipitates appeared (Figure 3C). Based on FCS measurements (Figure S5B), the fraction of protein incorporated into these aggregates is very low. Still, this indicates that interactions between specific RNA and the RRMs can lead to protein-RNA co-assembly. Such interactions with U-20 are likely too weak to lead to co-assembly at the same concentration.

Altogether, these results indicate that hnRNPA1-RNA cluster formation results from the multi-domain architecture of hnRNPA1 and requires both folded and intrinsically disordered domains.

### Characterization of hnRNPA1-RNA clusters

We next sought to unravel the different properties of the protein-RNA clusters observed at high RNA concentration with respect to the condensates formed at lower stoichiometry. We first analyzed the stability of the clusters upon addition of RNAse A. As indicated by diffusivity changes of hnRNPA1 and RNA (Figure 4A), the clusters disassemble, demonstrating that intact RNA is necessary for cluster formation. Next, we assessed the role of electrostatic interactions in cluster formation by increasing the salt concentration to 250 mM NaCl. FCS diffusivity measurements showed that clusters disassemble into monomeric protein and RNA (Figure 4B), indicating that electrostatic interactions are crucial for the formation of the hnRNPA1-RNA clusters similar to condensates, which do not form at this salt concentration.^5^

**Figure 4:**
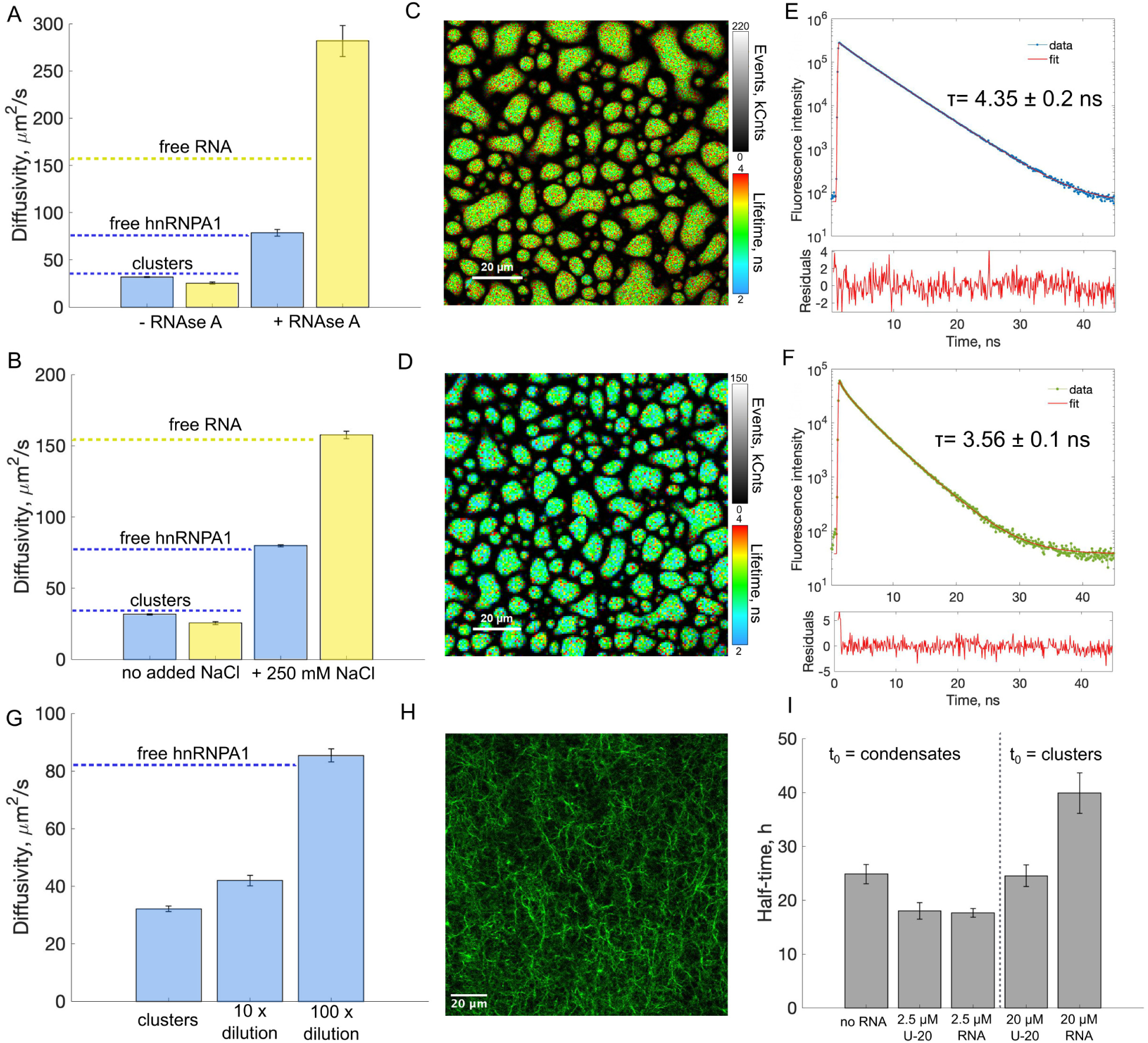
Characterization of hnRNPA1-RNA clusters formed at high RNA concentration. A-B) Clusters disassemble upon treatment with RNAse A (A) and upon addition of 250 mM NaCl (B). C) Atto-647 FLIM image of condensates formed in a sample containing 10 *µ*M hnRNPA1 (incl. 500 nM hnRNPA1-647) and 2.5 *µ*M RNA (incl. 500 nM RNA-FAM). D) Fluorescein (FAM) FLIM image of condensates formed in a sample containing 10 *µ*M hnRNPA1 (incl. 500 nM hnRNPA1-647) and 2.5 *µ*M RNA (incl. 500 nM RNA-FAM). E-F) Fluorescence lifetime of hnRNPA1-Atto-647 (E) and RNA-FAM (F) in clusters (10 *µ*M hnRNPA1 and 10 *µ*M RNA) measured with fluorescence lifetime correlation spectroscopy. The average lifetime of hnRNPA1-Atto-647 in condensates and clusters is 3.1 ± 0.2 ns and 4.35 ± 0.2 ns, respectively. The average lifetime of RNA-FAM in condensates and clusters is 2.7 ± 0.3 ns and 3.56 ± 0.1 ns, respectively. G) hnRNPA1-RNA clusters are unstable upon dilution as measured by diffusivity changes with FCS. H) hnRNPA1-RNA clusters are unstable over time and transition into amyloid fibrils. Fluorescence image of fibrils formed in a sample containing 10 *µ*M hnRNPA1, 10 *µ*M RNA and the amyloid reporter dye Proteostat incubated for 90 h. I) Half-times of aggregation kinetics traces measured in samples containing 10 *µ*M hnRNPA1 with no RNA, 2.5 *µ*M U-20 or RNA (two-phase regime) and 20 *µ*M U-20 or RNA (one-phase regime).

We also applied fluorescence lifetime measurements to compare the chemical environment of hnRNPA1-RNA condensates and clusters. Fluorescence lifetime microscopy (FLIM) images of hnRNPA1-647 and RNA-FAM in condensates are showed in Figure 4C and D, respectively. In condensates, fluorescence lifetime of Atto-647 conjugated to hnRNPA1 is 3.1 ± 0.2 ns, while the lifetime of FAM conjugated to RNA is 2.7 ± 0.3 ns. Fluorescence life-times inside the clusters were measured with Fluorescence Lifetime Correlation Spectroscopy (FLCS) (Figure 4E and F). The values are significantly larger than in the condensates and equal 4.35 ± 0.2 ns and 3.56 ± 0.1 ns for hnRNPA1-647 and RNA-FAM. These lifetimes are close to lifetimes measured for monomeric hnRNPA1-647 and RNA-FAM in buffer, which equal 4.3 ± 0.1 ns and 3.85 ± 0.05 ns, respectively (Figure S6). This might be explained by lower macromolecular density inside the clusters and/or higher exposure of the fluorescent dyes to the solvent compared to within the condensates. These data show that the chemical environment in the clusters is different from the one in the condensates and is close to the environment experienced by monomeric protein and RNA in solution.

Next, we investigated the stability of the clusters upon dilution (Figure 4G). 10-fold dilution of hnRNPA1-RNA clusters formed at 10 *µ*M protein and 10 *µ*M RNA leads to smaller clusters, which are 5 nm in radius compared to 7.3 nm before dilution. Upon further dilution (100x), the clusters immediately disassemble into monomeric protein and RNA.

Finally, we tested the stability of hnRNPA1-RNA clusters over time. Upon incubation for 90 hours, clusters convert into amyloid fibrils, as shown by confocal fluorescence microscopy (Figure 4H). This result indicates that the observed clusters are metastable species. Formation of amyloid fibrils was observed also in the regime where condensates are present, for both the LCD and the full length protein.^5,24,25^ When we compared the timescale for fibril formation in the two-phase regime with condensates and in the one-phase regime with protein-RNA clusters (Figure 4I and Figure S7), we observed that the transition to amyloid fibrils from hnRNPA1-RNA clusters is slower compared to condensates (aggregation halftime of (*t*_1_*_/_*_2_=39.9 ± 7.5 h versus *t*_1_*_/_*_2_=17.6 ± 1.7 h). We compared the effect of U-20 and RNA on the kinetics of amyloid formation. While no significant difference was observed in the two-phase regime (Figure 4I and Figure S7), aggregation was faster in the presence of U-20-protein clusters compared to RNA-protein clusters. These results show that clusters formed with RNA are kinetically more stable than those formed with U-20, suggesting that assemblies formed by proteins and specific RNAs can be more protective towards fibril formation compared to unspecific RNA molecules.

## Discussion

We have shown that interactions between hnRNPA1 and RNA can lead to two different types of assemblies, which are in competition with each other and are modulated by RNA concentration: nano-sized clusters and micron-sized condensates (Figure 5A). At low RNA concentration, hnRNPA1 phase separation is promoted, while at high RNA concentration phase separation is suppressed and soluble protein-RNA clusters form in the dilute phase. These results show that protein-RNA clusters represent a more efficient strategy to pack large amounts of RNA in protein-RNA assemblies than micron-sized condensates.

**Figure 5:**
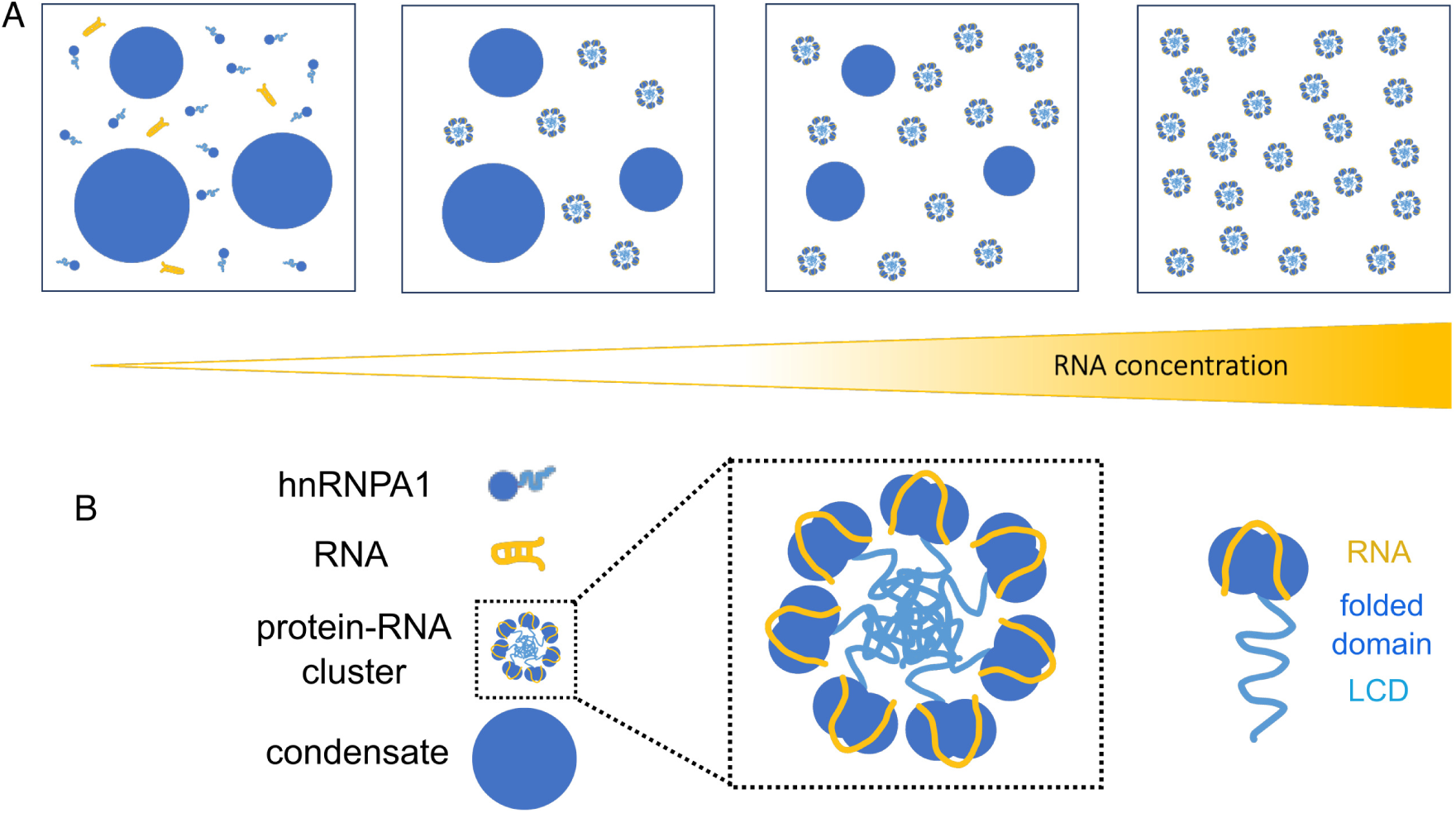
Competition with protein-RNA clustering in the dilute phase drives suppression of hnRNPA1 phase separation at high RNA concentration. A) Schematic illustration of the transitions of the system occuring at constant protein concentration with increasing RNA concentration. Within the investigated interval, there is a broad RNA concentration range where condensates and clusters in the dilute phase coexist. B) Hypothetical structure of the hnRNPA1-RNA cluster, where low complexity domains of the protein point towards the interior of the clusters, while folded domains to which RNA binds are exposed towards the exterior. The structure shown is a speculative representation based on available data and we cannot exclude that RNA can also be incorporated into the core of the structure.

Our results are consistent with and shed new light on the *in vivo* study of Maharana et al.,^4^ who showed that phase separation of several RNA-binding proteins, including hnRNPA1, is suppressed in the nucleus due to high RNA concentration. In-cell FCS experiments showed that RBPs in the nucleus exist predominantly as slowly-diffusing species,^4^ which in light of our results are likely protein-RNA clusters. Our results should also be discussed in the context of 40S hnRNP particles formed from hnRNP proteins and nuclear RNA.^26,27^ These particles were recently extracted from nuclei, shown to be approx. 10 nm in radius and composed mainly of hnRNPA1 and hnRNPC proteins. ^27^ Their formation depends on RNA as RNAse treatment led to their disassembly and requires the LCD of hnRNPA1, as similar particles were not observed for the folded domain only (UP1).^27^ Thus, the 40S hnRNP particles share many characteristics with the hnRNPA1-RNA clusters that we characterize in this work. They are likely functional complexes in the different mRNA processing steps that hnRNPA1 plays a role in.

A question that naturally emerges is how the hnRNPA1-RNA clusters differ from the condensates in terms of the underlying protein-protein and protein-RNA interactions. First, the characteristic structure of hnRNPA1 featuring a folded domain and a low complexity domain plays a crucial role in cluster formation. No protein-RNA clustering in the dilute phase was observed in the case of UP1 or for A1-LCD. Thus, both domains of hnRNPA1 are necessary for cluster formation, while the low complexity domain alone is necessary and sufficient for condensate formation.^16^

hnRNPA1 can engage in interactions with RNA through the RRMs in the folded domain and through the RGG repeats in the LCD.^15,28^ We expect that both types of interactions contribute to phase separation as well as cluster formation, but likely with different relative contributions. Our data show that 18nt SMN2-ISS-N1 RNA is more efficient in suppressing phase separation than U-20 (Figure 2), but they are equally effective in promoting phase separation. This important observation suggests that high-affinity and sequence-specific RNA interactions with the RRMs play a more important role in cluster than in condensate formation. Thus, hnRNPA1 phase behavior is not only modulated by RNA concentration but also its chemistry. This effect has potential implications in modulating phase behavior of hnRNPA1 *in vivo*, since the protein is exposed to distinct RNA sequences in different compartments, with which it can interact with a wide range of affinities. Indeed, in the case of FUS, Maharana et al. showed that the fraction of slow-moving proteins in the nucleus, which we interpret as FUS-RNA clusters, decreased by weakening the strength of protein-RNA interactions, which were modulated through mutations in the RNA-binding domains. ^4^ Moreover, FUS variants with a lower affinity for RNA were more prone to phase separation,^4^ which in light of our work is likely due to a decreased propensity to form soluble clusters. This interpretation could also explain the observations of Mann et al., who showed that phase separation of an RNA-binding deficient mutant of TDP-43 generated by introducing point mutations in the RRM (thus interfering with sequence-specific protein-RNA interactions) was not suppressed at high RNA concentration in contrast to wild type TDP-43.^7^ This is consistent with a decreased propensity of the mutant to form clusters with RNA.

Our findings may have important biological relevance in the context of protein aggregation associated with neurodegenerative diseases, where subcellular localization of RBPs plays an important role.^29–33^ For instance, in ALS, amyloid aggregates are found in the cytoplasm, but not in the nucleus.^34–37^ Moreover, disease-related mutations affect partitioning of the protein between nucleus and cytoplasm.^29,38^ The combined effect of different concentrations and identities of RNA in the two different cellular compartments likely modulates not only the physiological but also pathological functions of hnRNPA1. hnRNPA1 amyloid formation was already shown to be modulated by RNA in both the phase separation-promoting and suppressing regimes.^5^ This work shows that nano-sized protein-RNA clusters formed at high RNA concentration slow down amyloid formation when compared to condensates, consistent with the fact that hnRNPA1 amyloid fibrils are not found in the nucleus.^4^ This is consistent with previous findings on the protective role of TDP-43 oligomers against fibril formation.^39,40^ Our data show that RNA chemistry also plays a role: transition from protein-RNA clusters to amyloid fibrils is slower in the case of specific RNA when compared to U-20. This is likely due to higher stability of hnRNPA1-RNA clusters and suggests that their further stabilization might be a strategy to inhibit amyloid formation.

We propose a structure of hnRNPA1-RNA clusters by drawing an analogy to the structure of a charged surfactant micelle (Figure 5B). In a surfactant micelle, hydrophobic tails are sequestered inside, while headgroups are exposed outside. hnRNPA1 can be considered surfactant-like due to its multi-domain architecture with the highly charged folded domain acting as a headgroup and the low complexity domain acting as a tail. In a plausible structure of an hnRNPA1-RNA cluster, LCDs are oriented towards the interior, while folded domains to which RNA binds is located on the surface of the cluster (Figure 5B). The fact that clusters do not form in the absence of RNA despite large self-association propensity of the LCDs suggests that the intramolecular interactions between the different domains of hnRNPA1 are not repulsive enough for the micelle-like structure to be favored, consistent with computational predictions by Shinn et al.^41^ RNA binding perturbs the intramolecular interactions between the folded domain and LCD,^23^ which might render the molecule more prone to micelle formation. The effects of RNA and U-20 on cluster formation (Figure 2) support the micelle-like model of the clusters: we would not expect a large difference in the effect of these two RNA types on cluster formation if the RNAs in the clusters were interacting predominantly with the LCDs. Given that 18nt SMN2-ISS-N1 RNA interacts with high affinity with the RRMs of hnRNPA1,^15,23^ its higher efficiency in cluster formation likely reflects the mode of interaction in the clusters - namely, association with RRMs rather than with the LCD. Moreover, the proposed structure is consistent with the fact the clusters exhibit values of fluorescence lifetime similar to monomeric protein and RNA (Figure 4E,F and Figure S6).

Beyond RNA-binding proteins associated with neuro-degeneration, competition between (stoichiometric) clustering and phase separation is emerging as a potential more general mechanism by which cells can modulate assembly of macromolecules.^42–44^ For instance, Seim et al. demonstrated that the phenomenon governs phase behavior of a fungal ribonucleoprotein Whi3 both in absence and presence of RNA.^42^ Competition between self-assembly and phase separation was also proposed to modulate phase behavior of an enzyme Rubisco and a protein EPYC1, components of pyrenoid - a membraneless organelle involved in carbon fixation.^43^

## Conclusions

We have shown that dissolution of full-length hnRNPA1 condensates with increasing concentration of RNA is driven by a competition with the formation of nano-sized protein-RNA clusters in the dilute phase. This mechanism fundamentally differs from the charge inversion effect that induces dissolution of complex coacervates when increasing the concentration of one charged component.

Competition between clustering and phase separation requires both domains of hnRNPA1 and is modulated not only by RNA concentration but also by the strength of protein-RNA interactions: specific RNA is more effective in promoting cluster formation than unspecific RNA, although their effect in inducing condensates at low concentrations is similar. Protein-RNA clusters are metastable and convert over time into amyloid fibrils, but over a longer time-scale compared to conditions with condensates.

The competition between clustering and phase separation reported in this study provides a unifying framework to understand the different assemblies of hnRNPA1 in different cellular compartments, with potential implications for its physiological and pathological functions. The fact that protein-RNA cluster formation suppresses phase separation and delays amyloid fibril formation is consistent with the reported absence of aggregation in the nucleus.^4^ In the cytoplasm, vice-versa, non-sequence-specific interactions with RNA present at lower concentrations might induce phase separation of hnRNPA1 rather than cluster formation, which in addition could be less effective in preventing fibril formation over time.

## Materials and Methods

### Protein purification and fluorescent labeling

Full length hnRNPA1A and UP1 was expressed and purified as described previously.^5^ Labeling with Atto 647N NHS ester dye (Atto-Tec GmbH, Germany) was done by purifying in 10 mM phosphate buffer at pH 7.4 containing 500 mM NaCl and incubating for 2 h at room temperature with the dye at 1:1 dye:protein molar ratio. After incubation with the dye the sample was loaded on Superdex 75 column (Cytiva Sweden AB, Sweden) equilibrated with 50 mM TRIS at pH 7.4 containing 500 mM NaCl and 10% glycerol. The collected sample was concentrated to 100-200 *µ*M, aliquoted into 5 *µ*l and flash-frozen in liquid nitrogen. The hnRNPA1A-LCD was purified as described previously.^24^ For fluorescent labeling with Atto 647N NHS ester, the protein was incubated with 1:1 protein:dye molar ratio overnight. Before gel filtration, the sample was centrifuged to remove any aggregates formed during incubation.

### RNA

Single-stranded RNA with a sequence is 5*^′^*-CCA GCA UUA UGA AAG UGA-3*^′^* and the same sequence labelled at the 3*^′^* end with carboxyfluorescein (FAM) as well as homopolymeric RNA U-20 and U-20 labelled at 3*^′^* end with carboxyfluorescein (FAM) were purchased from Microsynth AG (Baglach, Switzerland) as HPLC-purified lyophilized solids and were used without further purification.

### Sample preparation

All the experiments (apart from turbidity measurements) were performed using glass bottom 384-well plates (Azenta Life Sciences). Condensation was triggered by diluting protein stock 10 times in buffer containing 20 mM Tris at pH 7.5 and the final sample volume was 20 *µ*l.

### Right angle scattering measurements

Right angle scattering measurements were carried out with Labbot (Probation Labs Sweden AB). The sample in a quartz cuvette (Hellma GmbH & Co. KG, Germany) with a path length of 3 mm was illuminated with a 635 nm laser and scattered light intensity was measured at a 90*^◦^*angle. The values plotted in Figure 1C and Figure 2C are a mean of three biological replicates measured 5 min after inducing phase separation and normalized with respect to the maximum value in the concentration series.

### Dynamic Light Scattering

DLS experiments were performed using Prometheus Panta (NanoTemper GmbH, Munich, Germany). Single–use high sensitivity glass capillaries were filled with several *µ*l of the sample. Data analysis was performed using Prometheus Panta Control Software.

### Confocal imaging

Confocal imaging was performed on an inverted confocal fluorescence microscope (Leica SP8 STED, Leica Application Suite X (LAS X) software, version 1.0) equipped with a HC PL APO CS2 63x 1.2 NA water immersion objective and a hybrid detector for single molecule detection (HyD SMD).

### Fluorescence correlation spectroscopy

FCS experiments were performed on an inverted confocal fluorescence microscope (Leica SP8 STED, Leica Application Suite X (LAS X) software, version 1.0) equipped with a HC PL APO CS2 63x 1.2 NA water immersion objective with a software-controlled correction collar (Leica) and a hybrid detector for single molecule detection (HyD SMD). The confocal volume of the 488, 565 and 647 channels was calibrated using Atto 488 NHS Ester (*D_coeff_* = 400 *µm*^2^/s), Atto 565 NHS Ester (*D_coeff_* = 400 *µm*^2^/s) and Alexa 647 NHS Ester (*D_coeff_* = 330 *µm*^2^/s), which yielded an effective volume, *V_eff_*, of 0.3 ± 0.05 fl and a focal volume height-width ratio, *κ* = 6, *V_eff_*, of 0.4 ± 0.1 fl and *κ* = 6.5 and *V_eff_*, of 0.6 ± 0.1 fl and *κ* = 6, for the different channels respectively.

The samples were excited with a 488 nm, 565 nm or 633 nm laser (from a white Light Laser at 80 MHz) and the fluorescence emission collected in the range 500-530 nm, 570-600 nm and 650-700 nm, respectively. For the measurements carried out in the dilute phase of the two-phase systems, it was ensured that directly below the chosen spots for analysis there are no condensates on the surface of the well.

### Fluorescence lifetime correlation spectroscopy

FLCS experiments were carried out exactly as described in the case of standard FCS experiments, but the laser frequency used was 20 MHz. Fluorescence decay curves were fitted with a two-component reconvolution model and the values reported in text are the intensity-weighted average fluorescence lifetimes.

### Fluorescence lifetime imaging microscopy

FLIM experiments were performed on an inverted confocal fluorescence microscope (Leica SP8 STED, Leica Application Suite X (LAS X) software, version 1.0) equipped with a HC PL APO CS2 63x 1.2 NA water immersion objective with a software-controlled correction collar (Leica) and a hybrid detector for single molecule detection (HyD SMD). FLIM images of the droplets were acquired until 300 photons per pixel were collected in the brightest channel. The samples were excited with a 488 or 633 nm laser (from a White Light Laser at 20 MHz) and the fluorescence emission collected between 500 and 530 nm or 650 and 700 nm for the green and red channel, respectively. Data analysis was performed using SymPhoTime 64 2.1 software (PicoQuant, Berlin, Germany). Images were fitted pixel by pixel with a one component reconvolution model.

### RNAse A experiment

20 *µ*g/ml RNAse A stock solution was prepared in 20 mM TRIS buffer at pH 7.5. 1 *µ*l of this solution was added to 20 *µ*l of sample containing 10 *µ*M hnRNPA1A (incl. 1 *µ*M hnRNPA1A-647) and 10 *µ*M RNA (corresponding to 0.6 *µ*g/ml) to reach 1:100 RNAse:RNA molar ratio. FCS experiments were carried out before and 15 min after RNAse A addition.

### Aggregation kinetics

To monitor fibril assembly, we used the dye Proteostat (Enzo Life Sciences AG, Switzerland), because it shows reduced background signal in presence of ribonucleic acids compared to Thioflavin T. The dye stock was diluted 1000 times in the aggregation buffer (20 mM TRIS pH 7.5) at the beginning of the reaction. The increase in the fluorescence signal over time was monitored every 30 minutes on a ClarioStar plate reader (BMG Labtech, Germany) using the Reader Control and MARS software. The excitation light was set to 550 nm, while fluorescence emission was recorded at 600 nm.

## Acknowledgements

We kindly acknowledge the European Research Council through the Horizon 2020 research and innovation programme for financial support (grant agreement No. 101002094). We would like to thank Felix Roosen-Runge from Lund University and Jannette Carey from Princeton University for helpful discussions.

## Author contributions

K.M. and P.A. designed the study. K.M. carried out the experimental work and data analysis with the help of C.M. C.M. and L.F. contributed with protein expression and purification.

K.M. wrote the manuscript with input from all co-authors. P.A. supervised the project and acquired funding.

