## Supplementary information for "Competition between protein-RNA clustering and phase separation drives re-entrant phase behavior of hnRNPA1"

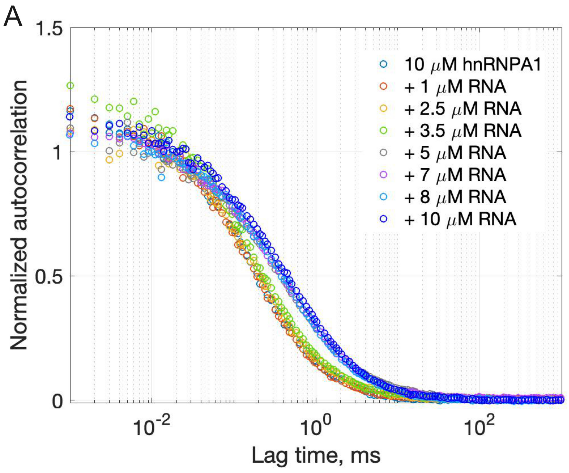


Figure S1: FCS autocorrelation curves of hnRNPA1-647 (A) and RNA-FAM (B) measured in the dilute phase of samples containing 10 $\mu$M hnRNPA1 (including 500 nM hnRNPA1-647) and increasing concentration of RNA (including 500 nM hnRNPA1-FAM). The shift towards longer lag times indicates formation of larger protein-RNA clusters in the dilute phase with increasing RNA concentration.

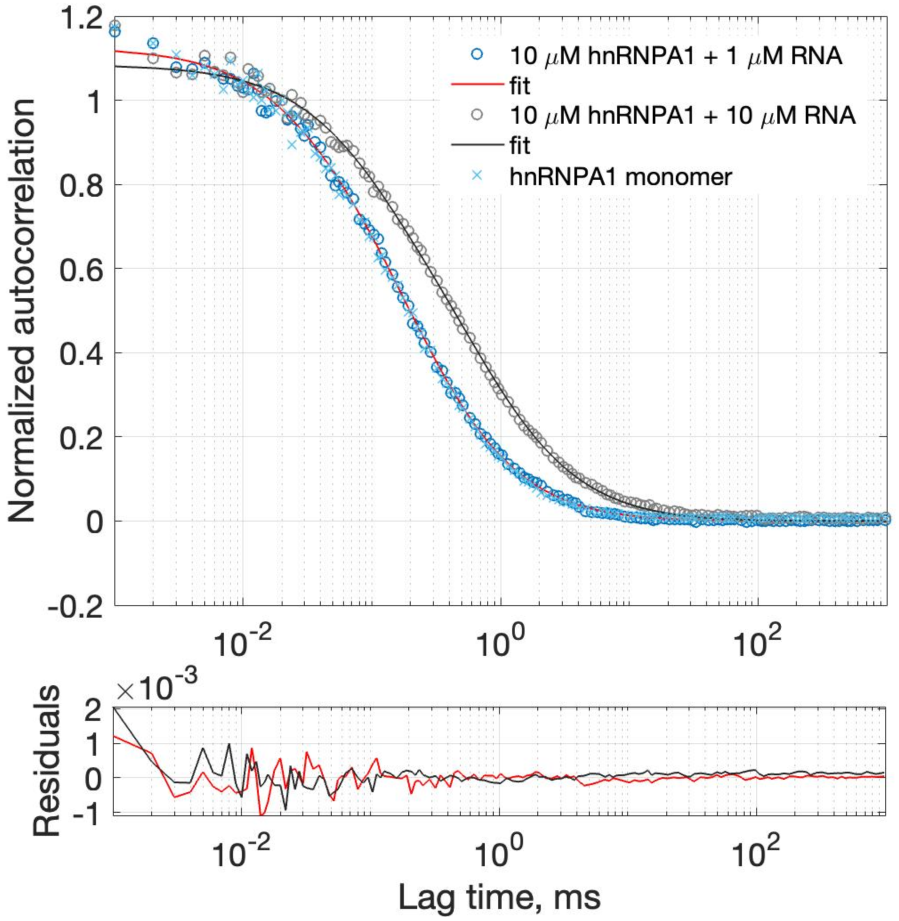


Figure S2: Fitting of hnRNPA1-647 FCS autocorrelation curves. The FCS autocorrelation curve of hnRNPA1-647 measured in the dilute phase coexisting with condensates at low RNA concentration (1 $\mu$M RNA, blue circles) overlaps with the autocorrelation curve of monomeric hnRNPA1 measured in buffer containing 500 mM NaCl (blue crosses). These curves could be fitted to a model assuming one diffusing component (diffusivity 82.3 $\pm$ 3.6 $\mu$m$^2$/s) and a triplet contribution with triplet time in the range 5-10 $\mu$s (red line). The FCS autocorrelation curve of hnRNPA1-647 measured in the one-phase regime at high RNA concentration (10 $\mu$M RNA, grey circles) also could be fitted to a model assuming one diffusing component (diffusivity 30.5 $\pm$ 0.7 $\mu$m$^2$/s) and a triplet contribution (black line). In this case, the triplet time fitted to 30-50 $\mu$s, which is too long to be due solely to triplet dynamics. This timescale is also too short to be due to diffusion. Having observed this fast contribution only in the samples containing protein-RNA clusters, we conclude that it arises from a process connected to the presence of clusters e.g., a protein conformational change or cluster assembly-disassembly dynamics. Here, we choose to account for this process in the fitting with a long triplet time. The accurateness of this approach is supported by the consistency of the FCS and DLS results (Figure 1E): at 10 $\mu$M hnRNPA1 and 10 $\mu$M RNA, both techniques show the presence of clusters of approx. 10 nm in radius and absence of monomeric protein.

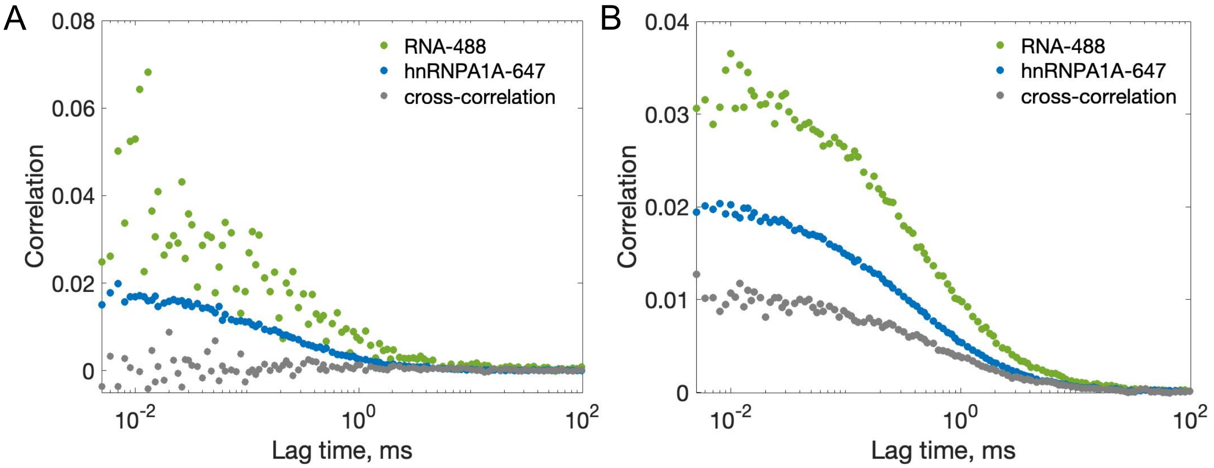


Figure S3: hnRNPA1A and RNA are co-assembled only in the dilute phase at high RNA concentration. A) Fluorescence cross-correlation experiment in the dilute phase of sample containing 10 $\mu$M hnRNPA1A and 1.25 $\mu$M RNA. The zero amplitude of the cross-correlation function (grey) indicates the absence of protein-RNA species in the sample. B) Fluorescence cross-correlation experiment in the sample containing 10 $\mu$M hnRNPA1A and 10 $\mu$M RNA. The cross-correlation curve reports on the dynamic co-localization of protein and RNA.

### Determination of the number of protein monomers per cluster from FCS data

Based on the measured diffusivities of hnRNPA1 monomer and clusters, we can approximately estimate the number of protein monomers per cluster. The estimate is based on the scaling of diffusivity ($D$) with molecular weight ($M_w$) as:

$$D \propto {M_w}^{-1/3}, \tag{1}$$

and assuming that both monomer and clusters are spherical and have the same relative density.[1] Thus, for diffusivity of hnRNPA1 monomer of 80 $\mu$m$^2$/s and hnRNPA1 cluster of 30 $\mu$m$^2$/s we get:

$$\frac{M_{w,cluster}}{M_{w,monomer}} = (\frac{D_{monomer}}{D_{cluster}})^3 \propto 20 \tag{2}$$

where $M_{w,cluster}$ and $M_{w,monomer}$ are the molecular weights of hnRNPA1 cluster and monomer, respectively. In this estimate, the contribution of RNA to the molecular weight of the cluster is omitted as it is much lower than the molecular weight of the protein (34 kDa and 6 kDa for monomeric hnRNPA1 and RNA, respectively).

### Determination of cluster stoichiometry from FCS data

We extract the number of fluorescently-labeled protein and RNA per cluster, and from this the RNA:protein stoichiometry of the cluster according to the following procedure.

First, we measure the total number of labeled protein molecules ($n_{647tot}$) in reference sample containing 10 $\mu$M hnRNPA1 incl. 500 nM hnRNPA1-647 in 500 mM NaCl where no phase separation nor clustering occurs, and the number of labeled RNA molecules ($n_{488tot}$) in a reference sample containing 10 $\mu$M RNA incl. 500 nM RNA-FAM. Next, we measure the the number of labeled particles ($n_{488}$ and $n_{647}$) in samples containing 10 $\mu$M hnRNPA1 (incl. 500 nM hnRNPA1-657) and 10 $\mu$M RNA (incl. 500 nM RNA-FAM). These numbers correspond to the number of protein-RNA clusters detected in 488 and 647 channels, given that the sample consists entirely of protein-RNA clusters (Figure S2 and Figure 1E).

Next, we calculate the number of fluorescently-labeled protein/RNA per cluster from:

$$n_{protein/cluster} = \frac{n_{647tot}}{n_{647}} \tag{3}$$

and

$$n_{RNA/cluster} = \frac{n_{488tot}}{n_{488}} \tag{4}$$

This holds only if the number of labelled protein/RNA is not limiting the number of clusters that we detect. This applies in our case, since for all samples measured $n_{488tot} >> n_{488}$ and $n_{647tot} >> n_{647}$ (see Table S1).

Finally, we calculate the cluster stoichiometry from:

$$RNA : protein = \frac{n_{RNA/cluster}}{n_{protein/cluster}} \tag{5}$$

Table S1: Calculation of RNA:hnRNPA1 stoichiometry in the clusters from particle numbers extracted from Fluorescence Correlation Spectroscopy data.

| Sample | N ± SD (n=3) |
|---|---|
| 10 $\mu$M RNA incl. 500 nM RNA-FAM ($n_{488tot}$) | 131.9 ± 3.8 |
| 10 $\mu$M hnRNPA1 incl. 500 nM hnRNPA1-647 ($n_{647tot}$) | 93.2 ± 2.5 |
| RNA-FAM in 10 $\mu$M hnRNPA1 and 10 $\mu$M RNA ($n_{488}$) | 87.3 ± 2.5 |
| hnRNPA1-647 in 10 $\mu$M hnRNPA1 and 10 $\mu$M RNA ($n_{647}$) | 69.1 ± 1.4 |
| number of labelled RNA per cluster | 1.5 ± 0.1 |
| number of labelled protein per cluster | 1.35 ± 0.05 |
| RNA:protein in cluster | 1.12 ± 0.003 |

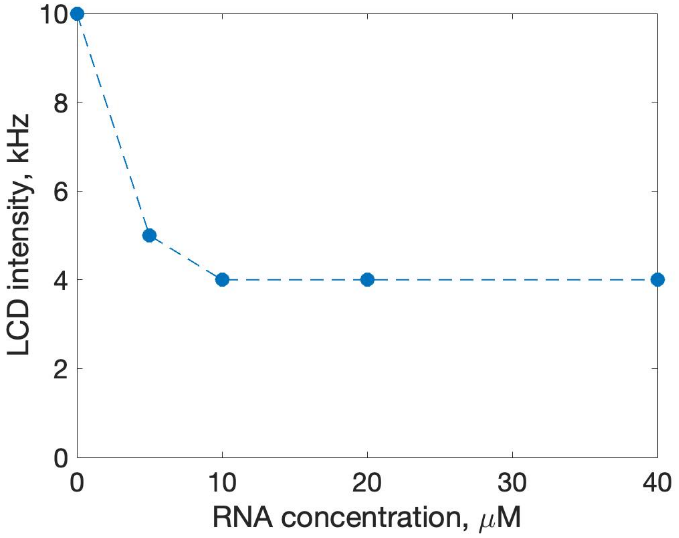


Figure S4: Fluorescence intensity of A1-LCD labelled with Atto 647 in the dilute phase of samples containing 10 $\mu$M A1-LCD and increasing concentration of RNA measured with Fluorescence Correlation Spectroscopy. The error bars are smaller than data points.

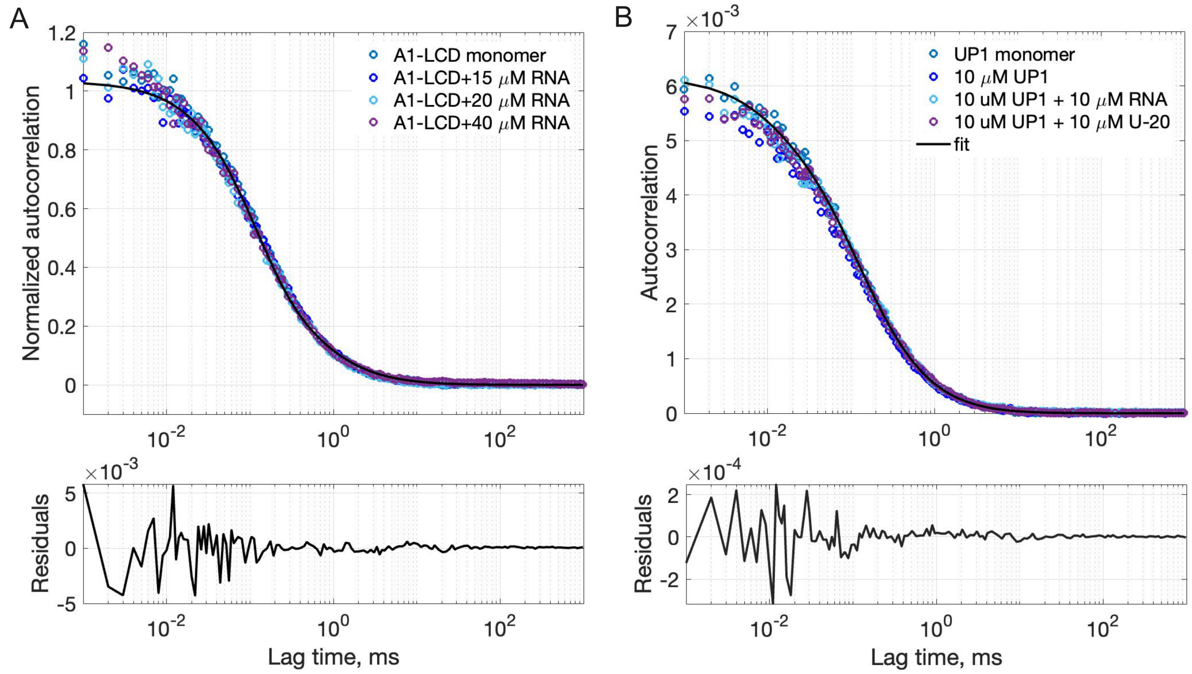


Figure S5: Diffusivity of A1-LCD and UP1 under different conditions studied with Fluorescence Correlation Spectroscopy. A) Autocorrelation curves of A1-LCD-647 in the dilute phase coexisting with condensates in absence and presence of 15, 20 and 40 $\mu$M RNA. The curves overlap with an autocorrelation curve of a monomeric A1-LCD and could be fitted to a model accounting for one diffusing component (fitted diffusivity 121.1 $\pm$ 2.8 $\mu m^2/s$) and a triplet contribution. B) Autocorrelation curves of 10 $\mu$M UP1 in absence and presence of 10 $\mu$M RNA or U-20. The curves overlap with an autocorrelation curve of a monomeric UP1 and could be fitted to a model accounting for one diffusing component (fitted diffusivity 94.5 $\pm$ 2.2 $\mu m^2/s$) and a triplet contribution. The almost identical amplitudes of all curves indicate that the aggregates observed in the sample containing RNA (Figure 3C) contain a negligible amount of total protein in the sample.

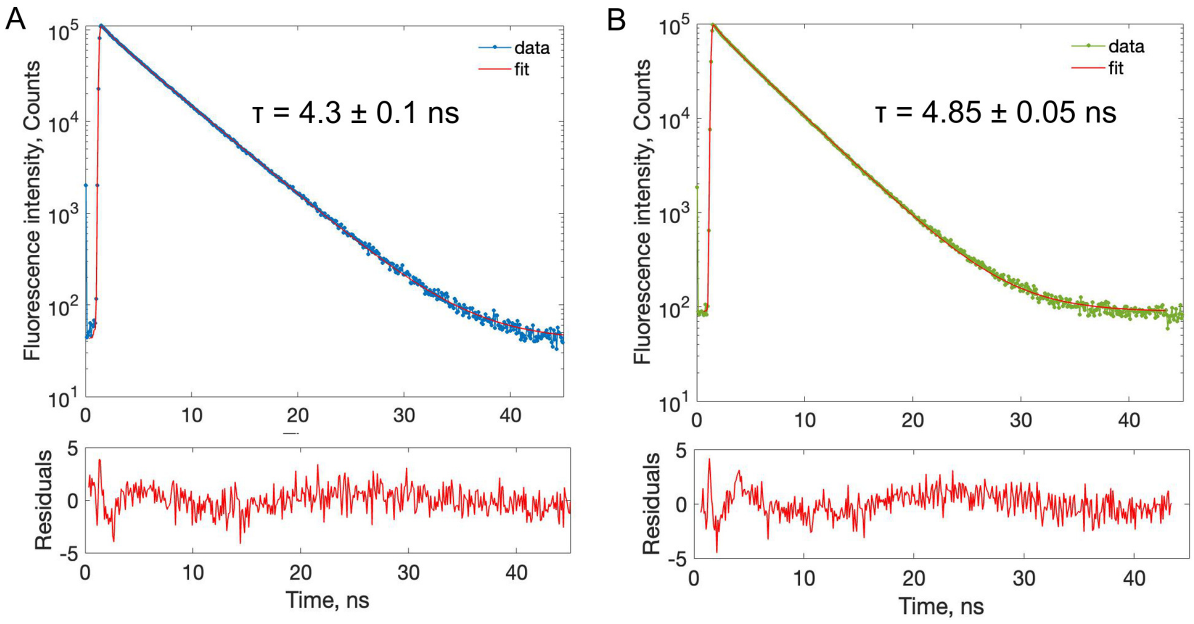


Figure S6: Fluorescence lifetime of Atto 647 conjugated to monomeric hnRNPA1 (A) and fluorescein (FAM) conjugated to RNA (B) measured in 20 mM TRIS buffer pH 7.5 with Fluorescence Lifetime Correlation Spectroscopy. Fluorescence decay curves were fitted with a two-component reconvolution model and the values reported are the intensity-weighted average fluorescence lifetimes.

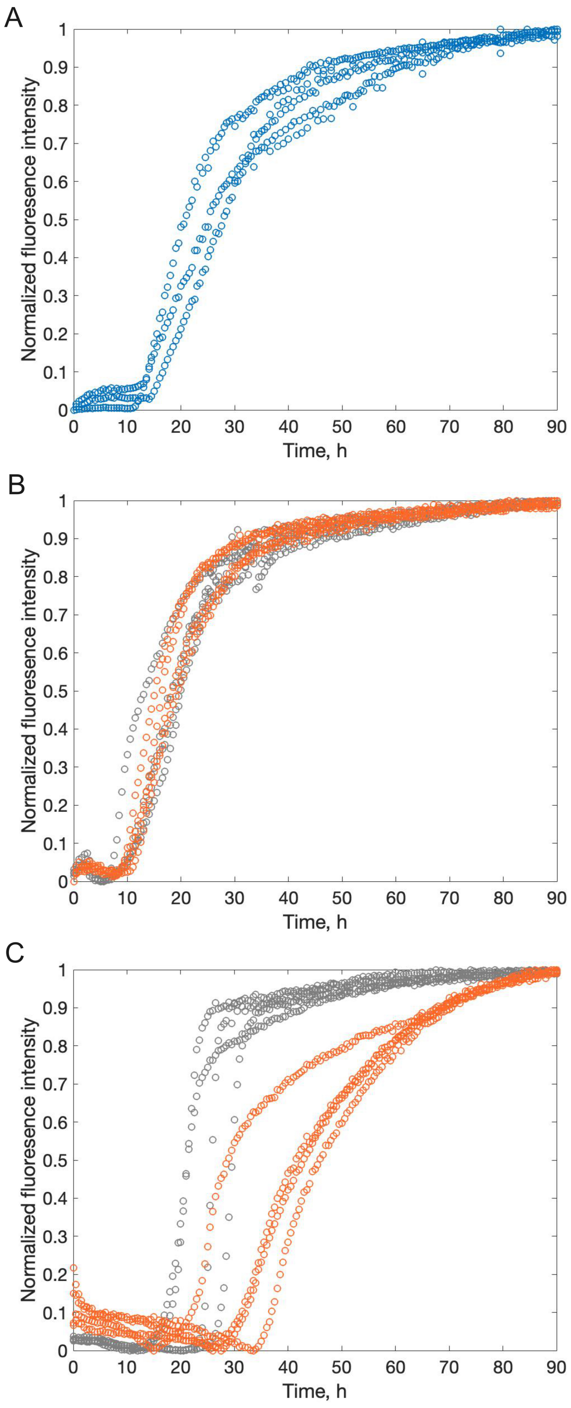


Figure S7: hnRNPA1 amyloid formation in the presence and absence of RNA. Aggregation kinetics of 10 $\mu$M hnRNPA1 in the two-phase regime with condensates in absence (A) and presence of 2.5 $\mu$M U-20 (grey circles) or RNA (orange circles) (B) and in the one-phase regime with protein-RNA clusters at 20 $\mu$M U-20 (grey circles)/RNA (orange circles)(C).
